## Supplemental Data for "In Vitro Evaluation and Mitigation of Niclosamide’s Liabilities as a COVID-19 Treatment"

### Supplemental Figures

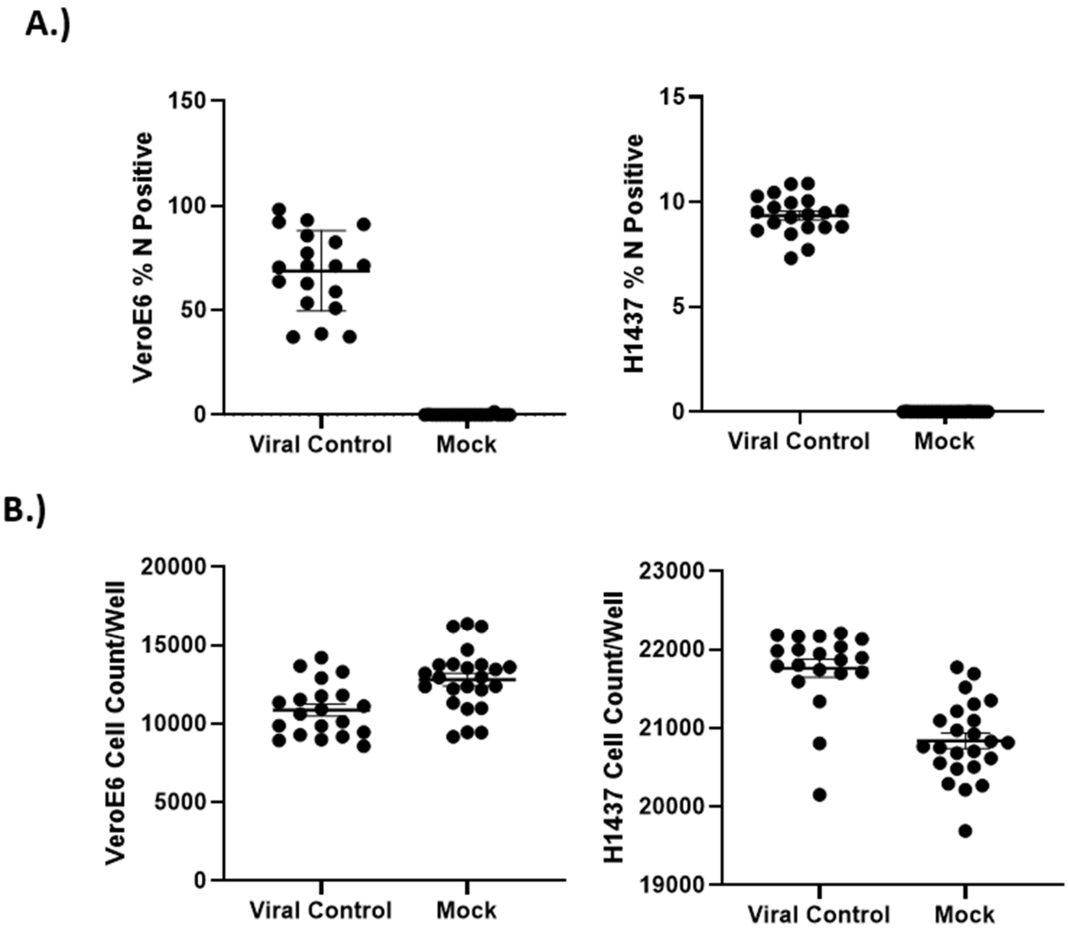


**Supplemental Figure S1.** Control data for VeroE6 and H1437 SARS-CoV-2 long-term exposure bioassays. Data were analyzed in GraphPad Prism 9.0 and is shown as mean ± standard deviation for N=20 replicates per condition. A.) Raw percent N protein positivity. Average percent infection was 69% for VeroE6 and 9% for H1437. This data was used to determine a normalized percent infection for treated wells. B.) Cell observations per well. Viral control data was used to determine a percent viability score for treated wells.

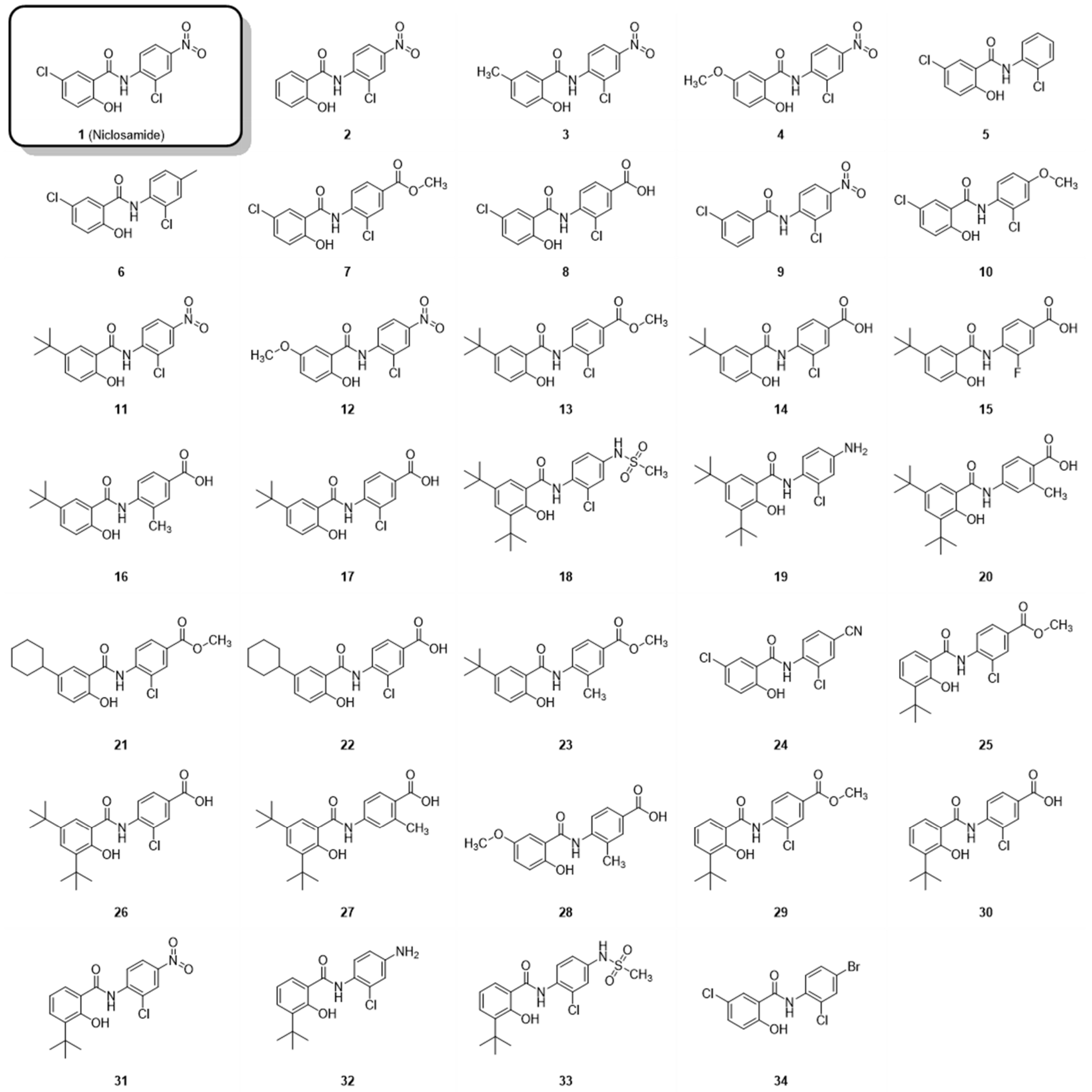


**Supplementary Figure S2.** Niclosamide analog structures. Modifications were made to the chlorosalicylic acid and/or nitroaniline rings. Compound number is indicated below the structure. Structures were generated using ChemDraw version 21.0.

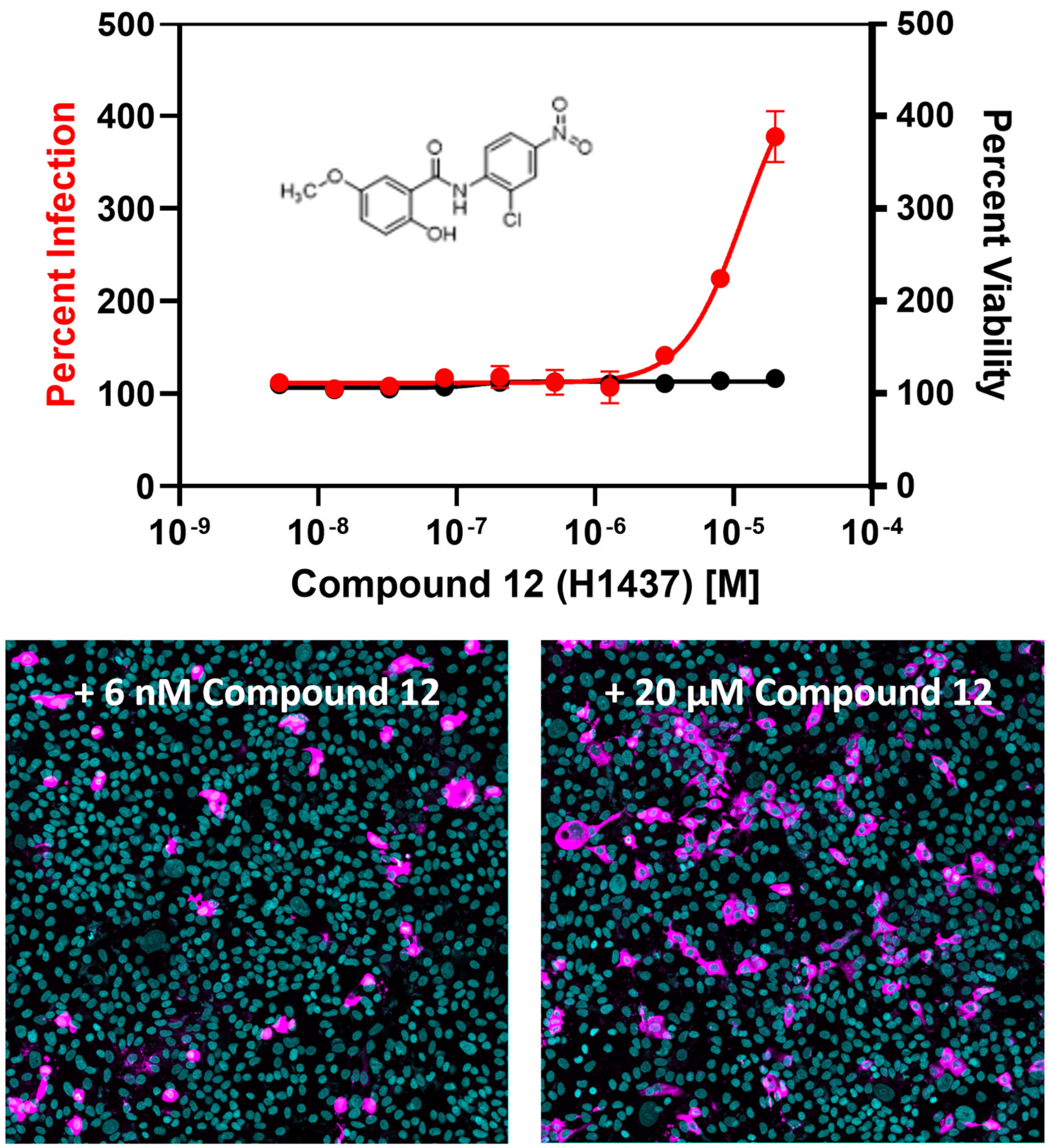


**Supplemental Figure S3. Some niclosamide analogs cause dose-dependent exacerbation of infection in H1437.** As a representative example, compound 12 causes a 400% increase in infection at 20 µM. A concentration-response curve for infection is shown here along with representative images for a non-exacerbating concentration (6 nM) and an exacerbating concentration (20 µM). Images were captured at 20X magnification using a Yokogawa CV8000 high content imaging microscope and processed using ImageJ. Nuclei are colored in cyan and SARS-CoV-2 N protein is colored in magenta.
